# Generalising Better: Applying Deep Learning to Integrate Deleteriousness Prediction Scores for Whole-Exome SNV Studies

**DOI:** 10.1101/126532

**Authors:** Ilia Korvigo, Andrey Afanasyev, Nikolay Romashchenko, Mihail Skoblov

## Abstract

Many automatic classifiers were introduced to aid inference of phenotypical effects of uncategorised nsSNVs (nonsynonymous Single Nucleotide Variations) in theoretical and medical applications. Lately, several meta-estimators have been proposed that combine different predictors, such as PolyPhen and SIFT, to integrate more information in a single score. Although many advances have been made in feature design and machine learning algorithms used, the shortage of high-quality reference data along with the bias towards intensively studied *in vitro* models call for improved generalisation ability in order to further increase classification accuracy and handle records with insufficient data. Since a meta-estimator basically combines different scoring systems with highly complicated nonlinear relationships, we investigated how deep learning (supervised and unsupervised), which is particularly efficient at discovering hierarchies of features, can improve classification performance. While it is believed that one should only use deep learning for high-dimensional input spaces and other models (logistic regression, support vector machines, Bayesian classifiers, etc) for simpler inputs, we still believe that the ability of neural networks to discover intricate structure in highly heterogenous datasets can aid a meta-estimator. We compare the performance with various popular predictors, many of which are recommended by the American College of Medical Genetics and Genomics (ACMG), as well as available deep learning-based predictors. Thanks to hardware acceleration we were able to use a computationally expensive genetic algorithm to stochastically optimise hyper-parameters over many generations. Overfitting was hindered by noise injection and dropout, limiting coadaptation of hidden units. Although we stress that this work was not conceived as a tool comparison, but rather an exploration of the possibilities of deep learning application in ensemble scores, our results show that even relatively simple modern neural networks can significantly improve both prediction accuracy and coverage. We provide open-access to our finest model at *http://score.generesearch.ru*.

## Introduction

Single amino-acid variation (caused by nonsynonymous single nucleotide substitutions – nsSNVs) is a valuable source of information that can help us understand the fundamental features of protein evolution and function as well as uncover causative variants behind inherent health conditions and develop custom treatment strategies to maximise therapeutic efficiency. The dramatic increase in our capacity to cheaply sequence human exomes has brought enormous amounts of information on genetic variation in human populations, which clearly has great potential in both theoretical and medical applications. The later fuels research towards the integration of personal genetic data into medical practice. In fact, various companies are already pushing the technology into consumer market, though the means to simplify and streamline the downstream analyses are still in the infancy, and our ability to interpret variation in a phenotypically-sensible manner leaves a lot to be desired. Untangling the connections between variation and phenotypic traits remains the greatest challenge of functional genomics, because only a small fraction of possible variants have been thoroughly investigated and manually reviewed with respect to their fitness impact, leaving the effects of most variants in the dark even when monogenic disorders are at stake. Thus, a lot of effort has been put into developing the means to infer possible damage of uncategorised nsSNVs by employing machine-learning. As a result, over the past decade many algorithms have been developed for predicting deleteriousness. In order to make predictions these tools encode variants using multiple quantitative and qualitative features, e.g. sequence homology [2], protein structure [3, 4] and evolutionary conservation [5, 6]. This diversity of scoring tools has led to the creation of dbNSFP [7–9], a regularly updated specialised database that accumulates predictions of various scores alongside genomic features for most of the possible variants in the human exome.

Meanwhile, the American College of Medical Genetics and Genomics (ACMG) published a guideline for reporting on clinical exomes [10], listing FATHMM [11], MutationAssessor [12], PANTHER [13], PhD-SNP [14], SIFT [15], SNPs&GO [16], MutationTaster [17], MutPred [18], PolyPhen-2 [19], PROVEAN [20], Condel [12], CADD [21], GERP [22], PhyloP [23] and several other scores as the most trustworthy. While the recommendations prove these scores useful, the guidelines describe them as merely accessory means of annotation, because differences in feature sets, training data and machine-learning algorithms used by the scores lead to inconsistent predictions (Fig 1), making the choice a matter of personal preference of each analyst [24].

**Fig 1.**
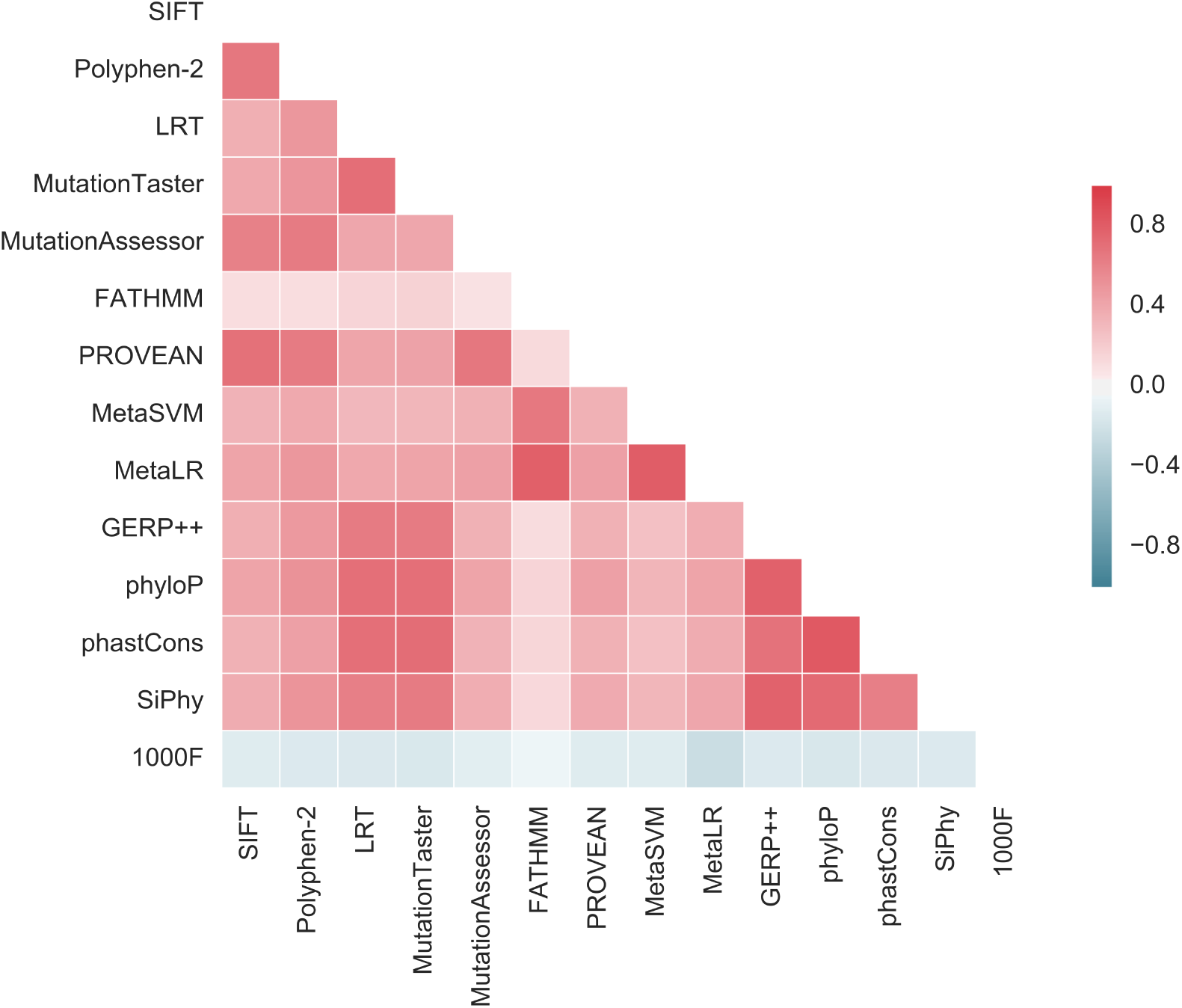
Prediction inconsistency. A heatmap of Spearman correlation between rank-transformed output values of different deleteriousness scoring systems. 1000F – allele frequency according to the 1000 Genomes project. Greater absolute correlation means greater consistency.

While several extensive comparison studies have been carried out [24–26], the differences in benchmarking datasets, the number of tools and precision assessment methods further complicate the generalisability of their conclusions. Therefore, it is still unclear which tools to use for prioritising variants in exome-based studies of human diseases. To reduce bias, gather more available information and simplify tool selection several meta-estimators have been proposed, based on other scores, such as PolyPhen and Sift. It has been demonstrated that combining individual predictions in ensembles results in mixed results. For example, KGGSeq (based on SIFT, PolyPhen-2, LRT, MutationTaster and PhyloP) [27] outperformed all its components in terms of ROC-curve AUC (area under the curve), while CONDEL (another meta-score) failed to do the same [24], though ROC-curves AUC are not the most comprehensive performance indicators. Following the trend, the curators of dbNSFP have developed their own ensemble scores (MetaLR and MetaSVM), that outperform all widely used standalone scores and meta-scores [24]. Additionally, to overcome the shortage of reference data, crucial in purely supervised training, some authors have proposed unsupervised and semi-supervised learning strategies, with CADD being the most notable implementation of the idea, though it doesn’t peform well in benchmarks [24]. Missing predictions (missing feature values) pose another serious problem. When one or more of the tools used by a meta-score fails to process a substitution (e.g. due to lacking some information about it) the entry becomes incomplete (Table 1) and thus requires special handling. Some tools handle missing values like an intrinsic property of the data [21], some try to impute them (by basically adding another machine learning task) [24], others are restricted to complete entries.

**Table 1.** Missing prediction rate of popular scores and meta-scores. *The fractions are estimated by querying a random subset of 1 * 10^6^ SNVs from dbNSFP v3.2 [9]. **Our testing datasets I and II (described in the Materials and Methods), comprising variations with experimental evidence of phenotype.

| Dataset | PolyPhen-2 | SIFT | FATHMM | MutationTaster | MetaLR |
| --- | --- | --- | --- | --- | --- |
| Exome* | 0.09 | 0.1 | 0.14 | 0.02 | 0.08 |
| Test I** | 0.02 | 0.03 | 0.06 | 0.01 | 0.004 |
| Test II** | 0.01 | 0.03 | 0.04 | 0.003 | 0.006 |

**Table 2.** ROC-curve AUC score.

| Score | Test I | Test II |
| --- | --- | --- |
| CADD | 0.85 (0.79-0.90) | 0.78 (0.78-0.79) |
| DANN | 0.84 (0.79-0.89) | 0.75 (0.74-0.75) |
| Eigen | 0.86 (0.81-0.91) | 0.67 (0.66-0.68) |
| FATHMM | 0.84 (0.78-0.89) | 0.91 (0.90-0.91) |
| GERP++ | 0.79 (0.73-0.84) | 0.68 (0.67-0.68) |
| LRT | 0.85 (0.80-0.90) | 0.73 (0.72-0.74) |
| **MLP** | 0.90 (0.86-0.94) | 0.94 (0.94-0.95) |
| MetaLR | 0.90 (0.86-0.94) | 0.93 (0.93-0.94) |
| MetaSVM | 0.90 (0.86-0.94) | 0.92 (0.92-0.93) |
| MutationAssessor | 0.78 (0.71-0.83) | 0.77 (0.77-0.78) |
| MutationTaster | 0.91 (0.87-0.94) | 0.76 (0.75-0.77) |
| PROVEAN | 0.82 (0.77-0.88) | 0.77 (0.77-0.78) |
| PolyPhen HDIV | 0.79 (0.73-0.85) | 0.77 (0.76-0.77) |
| PolyPhen HVAR | 0.80 (0.74-0.86) | 0.79 (0.78-0.79) |
| SIFT | 0.77 (0.71-0.82) | 0.78 (0.77-0.78) |
| SiPhy 29-way | 0.82 (0.76-0.87) | 0.70 (0.70-0.71) |
| phastCons 100-way | 0.81 (0.76-0.86) | 0.69 (0.69-0.70) |
| phyloP 100-way | 0.89 (0.85-0.93) | 0.75 (0.74-0.76) |
| phyloP 7-way | 0.80 (0.74-0.85) | 0.65 (0.64-0.66) |
| **sdAE** | 0.92 (0.88-0.95) | 0.92 (0.92-0.93) |

All these problems greatly emphasise the importance of better generalisation. Here we explore how deep learning can address the issues. Deep learning (DL) allows computational models learn hierarchies of representations with multiple levels of abstractions by combing several layers of nonlinear transformations. Techniques, such as noise injection and dropout, ultimately fight overfitting and allow adaptive regularisation [28]. Deep neural networks have already been used in DANN [1] and Eigen [29] to improve on the CADD’s original unsupervised approach, which incorporates hundreds of different features. While it is believed that one should only use DL for high-dimensional input spaces and other models (logistic regression, support vector machines, Bayesian classifiers, etc) for simpler inputs, we still believe that the ability of deep neural networks to discover intricate structure in highly heterogenous datasets can benefit a meta-estimator with relatively few input features, because connections and interaction between different scoring systems can be highly complicated and nonlinear [1]. We want to stress that this work was not conceived as a tool comparison, but rather an exploration of the possibilities of deep learning application in ensemble scores.

## Materials and Methods

### Testing and training data

Our testing setup is based on the extensive comparative study performed by Dong et al. [24]. Since MetaLR and MetaSVM, introduced in the study, were shown to be state of the art in meta-estimators, it was natural to include them here for the sake of comparison along with other scores evaluated in that study. Thus we had to make sure that our training and testing data did not give our models an unfair advantage, hence we used the testing datasets provided by the authors. Briefly, the authors constructed their first testing dataset out of 120 deleterious mutations (causing 49 different diseases) recently reported in Nature Genetics, and 124 neutral mutations newly discovered from the CHARGE sequencing project [30]. To ensure the quality of the deleterious mutations, they only left variants reported to cause Mendelian diseases with experimental evidence. The quality of the neutral mutations was ensured by removing any record with minor allele frequency < 1% in 2 thousands exomes from the ARIC study via the CHARGE sequencing project [30]. Additionally the authors used a subset of the nsSNV section of the VariBench dataset (the variants affecting protein tolerance) [26]. VariBench, comprising high quality records with experimentally verified effects, has become a standard dataset for performance evaluation. The subset included 6279 deleterious curated variants and 13240 common neutral variants (minor allele frequency > 1%).

UniProtKB/Swiss-Prot was the main source of annotated nsSNVs for our training dataset. We downloaded all the amino-acid natural variants (the HUMSAVAR archive) and mapped UniProt protein IDs to RefSeq nucleotide IDs. We then converted AA substitutions into nsSNVs. Initially there were around 28k causative and 39k neutral AA polymorphisms. We then downloaded ClinVar variants mapped to loci referenced in OMIM [31]. To remove untrustworthy deleterious variants, we manually read through 200 records and marked them as good or bad based on their descriptions. We then trained a bag-of-words Naïve Bayesian classifier to automatically process the remaining records. This mostly removed all records referencing various oncological conditions (we believe cancer-related variants, except for some rare cases, introduce nothing but noise to the dataset, because cancers are rarely monogenic and highly variable from the molecular point of view) or having nothing but *in silico* and/or GWAS-based evidence. This left us with around 120k variants. We further filtered them to remove any possible splicing-altering substitutions using the annotations from SnpEff 4.1 [32]. After that we removed records found in the testing datasets using the amino-acid notation. After all these steps there were around 96.5k variants left (64.5k deleterious and 32k neutral).

This was our raw supervised training dataset. For our final supervised training collection we only left true positive records with experimental evidence. The dataset comprised around 20k neutral and 15k damaging nsSNVs. For our unsupervised training dataset we simply sampled 10^6^ possible nsSNVs from the exome without replacement.

### Deep learning models

We constructed our classifiers using two basic architectures (Fig 2; a): the deep multilayer perceptron (MLP) and the stacked denoising autoencoder (sdAE). MLPs are well known and widely used models. Although their basic architecture was introduced decades ago, their modern versions differ significantly in many implementation details. Stacked denoising autoencoders are relatively novel models used for unsupervised and semi-supervised learning and data compression. These networks are first trained as individual shallow denoising autoencoders (Fig 2; b) by iteratively stacking one on top of another, which is followed by final training (Fig 2; c). The term ”denoising” stands for their ability to reconstruct lousy input records by generalising on training datasets. Stacking several autoencoders on top of each other and training each to reconstruct the output of the previous layer allows to learn a hierarchy of features in the input space in an unsupervised manner.

**Fig 2.**
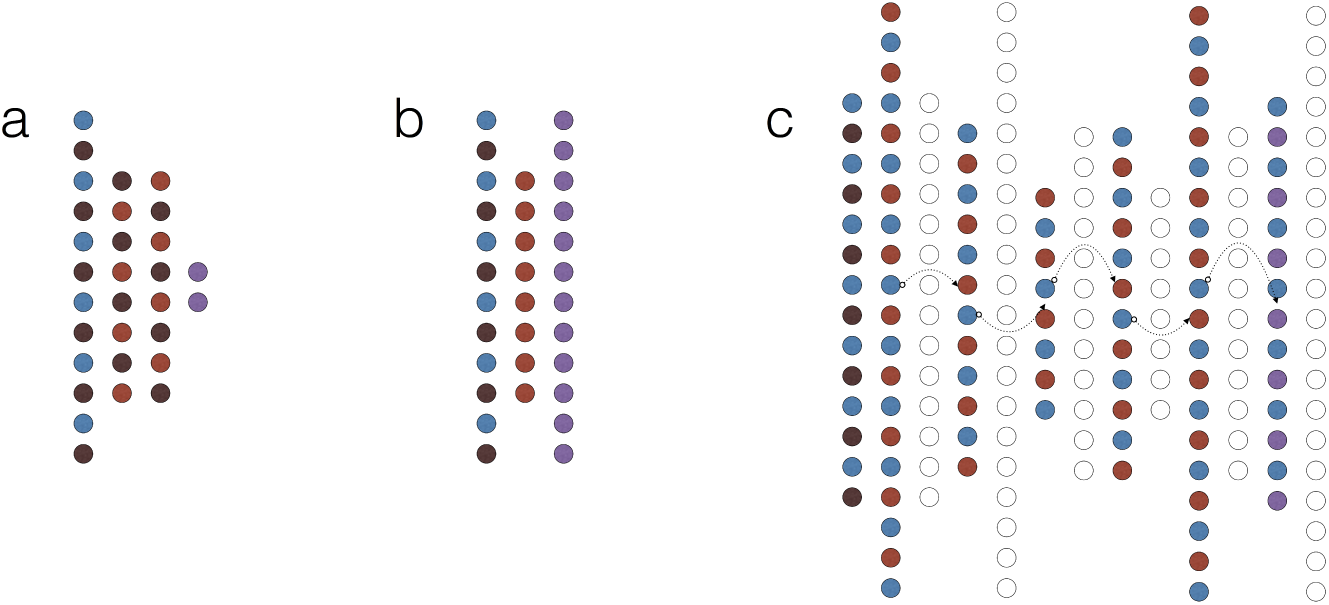
Network types. Schematic representation of basic deep learning models used in this study. (a) A multilayer perceptron (MLP). (b) A shallow denoising autoencoder (dAE). (c) Connecting dAEs into a stacked denoising autoencoder (sdAE); notice that each individual dAE learns to reconstruct the latent representation from the previous one (data stream is represented by arrows). Colours encode layer functions (combinations are possible): blue – input, light-red – latent, dark-red – dropout (noise), purple – output, hollow – discarded.

When labeled reference data are scarce, one can combine unsupervised and supervised training to discover great generalisations from unlabeled data and perform fine-tuning using the few available labeled records.

### Implementation details

Here we will briefly discuss several fundamental implementation details: update functions, regularisation, activation functions.

Most feed-forward networks train through stochatic gradient descent (SGD) combined with back-propagation of error. To make learning more efficient, many modifications of the regular stochastic gradient descent have been proposed. Here we explored two modifications: SGD with Nesterov momentum [33] and adagrad [34]. Both algorithms exploit the history of previous updates to prevent learning from slowing down and avoid local minima.

One of the greatest plagues of deep learning is actually a consequence of its very strengths – the multitude of neurons in several layers can coadapt to perfectly process the training dataset instead of learning valuable representations, which ultimately leads to overfitting. There are several ways to prevent this coadaptation and force a network to explore the feature space. Here we used dropout as a simple, yet extremely effective regularisation tool [28, 35]. During training, dropout can be interpreted as sampling a part within the full network, and only updating the parameters of the subsampled units during back-propagation. The process hinders coadaptation, because the subsampled networks share their parameters. In our MLPs we applied dropout to all layers, but the output, which can be interpreted as averaging predictions from an ensemble of networks. In sdAEs we only applied dropout to the input layer, which is often referred to as noise injection, which encourages autoencoders to discover intricate relationships between input units while trying to denoise them.

Activation functions are another highly important implementation detail. For our models we used sigmoidal and ReLU (rectified linear unit) nonlinearities. Briefly, the standard sigmoid function is defined as 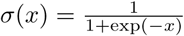, hence it maps any real number into (0, 1) and saturates at both ends, producing unfeasible gradients [36]. More importantly, repeated application of the sigmoid function (which basically happens when several hidden layers are used) leads to the vanishing gradient effect (Fig 3; a) hindering convergence. We also used the hyperbolic tangent (tanh), which is considered a superior sigmoidal function, because it is zero-centered and less prone to the vanishing gradient effect (Fig 3; b). The standard ReLU activation function is given by *ρ*(*x*) = *max*(0, *x*). Observe that this function has several nice properties:

- it has constant derivative defined as 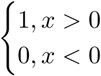
- it’s idempotent, that is its repeated application doesn’t affect the gradient, i.e. *ρ* o *ρ* o … o *ρ* = *ρ*
- it’s scaling invariant, i.e. *ρ*(*αx*) = *αρ*(*x*)
- it’s computationally cheap

These properties made ReLU the most popular nonlinearity of choice in most ANN architectures [37]. At the same time, it has been shown tricky to use the function in autoencoders, due to knockout effect and overshooting [38], hence it is still more common to use sigmoidal activations in these models.

### Hyper-parameter optimisation and the training setup

So far we’ve mentioned various aspects of design and implementation, influencing performance in many different ways [36]. These settings are called hyper-parameters: the number of layers and units per each layer, the compression factor in encoders, learning rate, dropout and noise levels, mini-batch size, momentum applied, nonlinearities used, etc. Usually, it takes a lot of time, experience and fine-tuning skills to come up with an optimal set of hyper-parameters. Alternatively one can try to optimise these using a special tool. For that we used a genetic algorithm (GA) – a stochastic optimisation algorithm simulating natural selection over many generations of mutation, recombination and selective pressure [39]. This strategy has already been successfully applied to optimise hyper-parameters in other machine-learning models [40]. Although powerful, genetic algorithms are extremely computationally expensive, especially in case of deep learning. Luckily, our models, due to the low-dimensional input space, required less weights resulting in faster convergence. At the same time, thanks to the advances in GPU-accelerated deep learning, the training time can be dramatically reduced when carried out on several GPUs [41]. We performed two independent GA runs: one for the pure MLP model and one for the stacked denoising autoencoder. In both cases a population of 100 individuals had been evolving for 100 generations and each model could chose whether to use the 7-way or 100-way phyloP and phastCons conservation scores, the batch size (500-10000), adagrad or Nesterov momentum (0.00-1.0; step size 0.05) update functions and the learning rate (0.005-1.0). During the MLP GA run the number of training epochs was fixed at 1000. All models used the hard ReLU activation function and Glorot uniform weight initialisation. The MLP-specific variable parameters:

**Fig 3.**
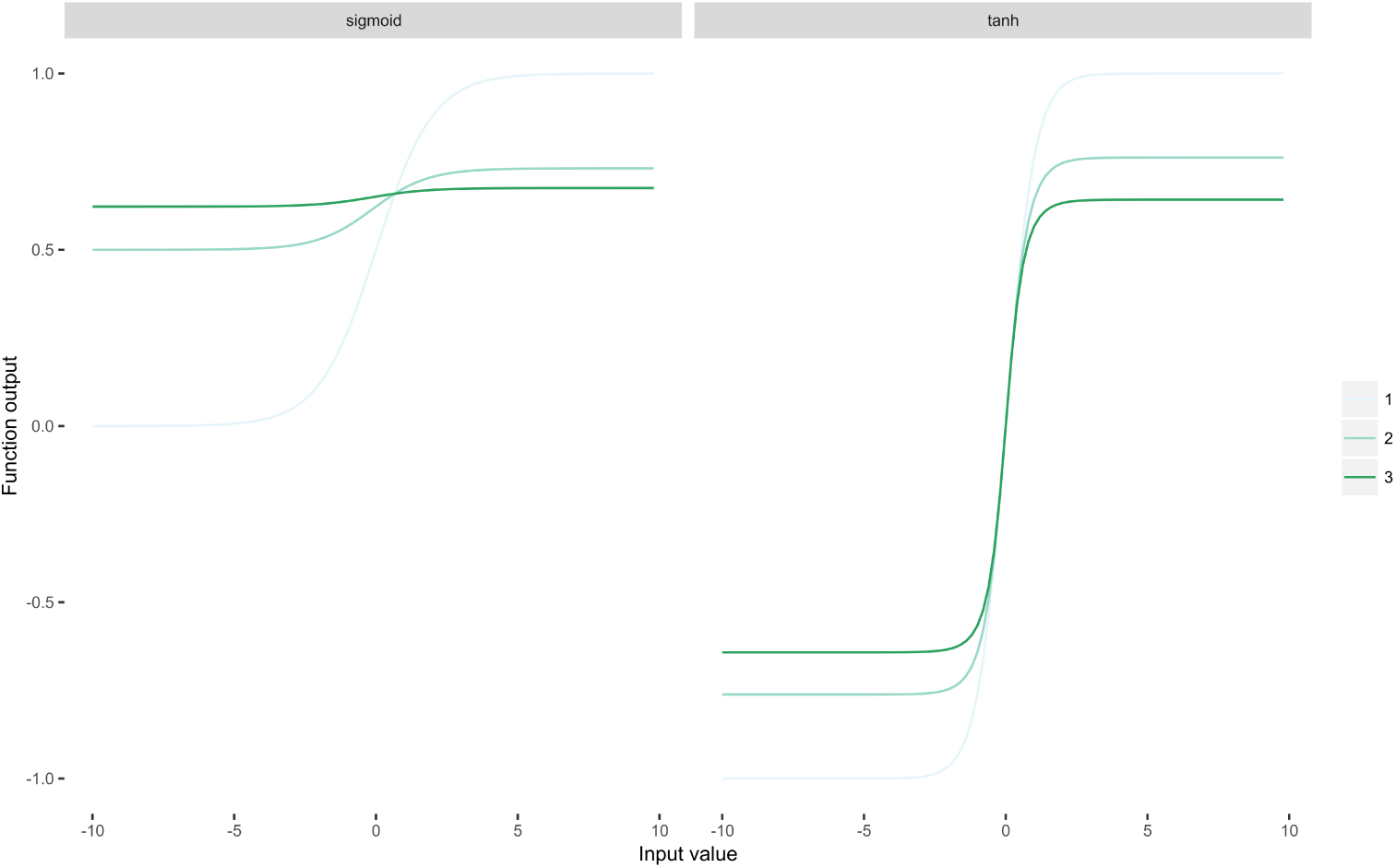
Nonlinearities. The sigmoid (a) and hyperbolic tangent (b) iteratively applied 3 times. Observe how repeated application of the sigmoid function quickly makes the gradient vanish completely.

- the number of hidden layers: 1-4
- the number of units per hidden layer: 10-30
- dropout probability: 0.00-0.5 (stepsize 0.05)

Each stacked denoising autoencoder trained in two steps: individual shallow autoencoders trained for 300 epochs prior stacking, then the stacked autoencoders trained for additional 1000 epochs. We increased the number of training epochs due to the saturation and vanishing gradient problems inherent to sigmoidal nonlinearities. The sdAE-specific variable parameters:

- first-layer expansion factor: 1.0-1.5 (stepsize 0.05); represents the relative increase in the number of units in the first hidden layer with respect to the input layer
- encoder compression level: 1.0-1.5 (stepsize 0.05)
- the number of hidden layers in the encoder (excluding the compressed latent layer): 1-3 (and the decoder by extension, due to symmetric design).
- activation function: sigmoid or hyperbolic tangent (in conjunction with appropriate weight initialisation functions).

We carried out the process on a machine with 8 Nvidia Titan X (Maxwell) GPUs using model-based parallelism, i.e. each model trained on a separate GPU with its own copy of the data, hence we could train up to 8 models simultaneously. The target function was the 3-fold cross-validation score (categorical crossentropy in case of the MLPs, and squared-root reconstruction error in case of the sdAEs). Neural network were implemented using Theano and lasagne in Python 3.5. We used the genetic algorithm implementation provided by package *genetic*, openly available in PyPI (the Python Package Index).

### Data extraction

We selected the following scores and genomic features as input units: FATHMM, GERP++, LRT, LRT Omega, MetaLR, MetaSVM, MutationAssessor, MutationTaster, PROVEAN, Polyphen2 (both HDIV and HVAR), SIFT, SiPhy log Odds, phastCons vertebrate (both 7-way and 100-way), phyloP vertebrate (both 7-way and 100-way) and the allele frequency (AF) in the 1000 Genomes Project dataset. Since all these scores had different output scales and thus couldn’t be directly compared, we used the rank-transformed values, provided by dbNSFP [7], for both training and comparison like demonstrated by Dong et al. [24]. Though from the training perspective this step was only mandatory for our autoencoders due to sigmoid activation in the decoders, other networks should’ve benefitted from the normalised bound data as well. For each position in the training and testing datasets we extracted these features from dbNSFP 3.1 and 3.2 [9] (we used the former to obtain values for the 7-way phyloP and phastCons, replaced by updated 100-way scores in the recent versions of dbNSFP).

### Handling missing data

We had no special way of handling missing data. Missing values in the training and testing datasets were simply replaced by zeros. We evaluated the performance of our classifiers without removing missing data from the dataset.

## Results and Discussion

### Training logs

We carried out two independent runs of genetic algorithm to optimise the hyper-parameters in our deep learning models. The MLP run took 3 days of calculations. We selected five sets of parameters yielding the highest cross-validation scores and trained them for 50k epochs. We then picked the network with the highest ROC-curve AUC and average precision (area under the precision-recall curve). The network had the following parameters:

- two latent layers: the first one had 13 hidden units, and the second one had 19
- dropout probability: 0.1
- batch size: 2000
- learning rate: 0.01
- update function: Nesterov momentum (0.7)
- phyloP and phastCons version: 7-way

The sdAE run took 54 days of calculations. As with the MLPs, we took 5 best-scoring models, though this time we trained each one for 100k epochs. After benchmarking the models on one million random nsSNVs from the exome (non-overlapping with the training dataset), we picked one model with the lowest absolute-error of reconstruction. It had the following parameters:

- three latent layers in the encoder
- expansion factor: 1.25
- compression factor: 1.3
- input-layer dropout (noise) probability: 0.3
- batch size: 5000
- learning rate: 0.05
- update function: Nesterov momentum (0.5)
- nonlinearity: hyperbolic tangent
- phyloP and phastCons version: 100-way

This model achieved median absolute reconstruction error of 0.02. We then removed the decoding part of the model, added a softmax output layer with two units and trained the model for 10k epochs to classify nsSNVs using the supervised training dataset. Training parameters were not altered, except for the batch size, which was reduced to 2000. Surprisingly, the resulting classifier performed poorly with average precision of 0.63 and ROC-curve AUC of 0.82, while having extremely low training errors, which led us to conclude that overfitting was the reason behind this results. To test this assumption, we tried to train the model again (staring with the same unmodified sdAE) while freezing the weights in the encoder and only updating the classifier’s softmax layer, which is basically similar to applying logistic regression on the compressed latent representation of the input space. This significantly increased both measures of performance. We used this final model in our benchmarks.

### Performance comparisons

The main performance indicators we used were the ROC-curve AUC score and the average accuracy score (area under the precision-recall curve), due to their threshold-invariance. We used bootstrapping to approximate the confidence intervals for these indicators. We also estimated the F1-score, Matthews correlation, accuracy, precision and recall on a range of binary cutoff thresholds (supplementary material S2). In our judgement we gave greater importance to the results of test II, because of the vastly greater size ( 100 times more records) and the origin (the Varibench benchmark) of the testing dataset II. The ROC-curve AUC tests supported the results published by Dong et al. [24] in their comparative study 2. The meta-estimators, introduced in that study (MetaLR and MetaSVM), outperformed most of the scores we used in the benchmark. Only our MLP classifier had a slight edge over both these scores in terms of the ROC-curve AUC. Though, MLP and MetaLR showed identical performance on the test I, which was second only to MutationTater and sdAE, the MLP outperformed all the other scores on the test II. At the same time the stacked autoencoder outperformed all scores on the test I. Surprisingly enough, the deep learning models that were developed to improve on the CADD’s unsupervised approach (DANN and Eigen) performed worse than CADD itself.

While ROC-curve AUC results gave our MLP a slight edge over the other scores on the test II, the average precision score deemed it even more superior 3, though it was second to sdAE on the test I. In general, most scores fared better on the testing dataset I in case of both performance indicators.

### Supervised vs. Unsupervised

Some researchers argue that unsupervised inference can help improve prediction quality, which can be hindered by poor coverage of the variome with reliable information on phenotypic categorisation. CADD is probably the most noticeable implementation of this approach. Since multiple studies, including this one, have shown that CADD performs significantly worse than most of its purely supervised rivals, it becomes unclear, whether there is any actual benefit of unsupervised learning in case of nsSNV classification. Most importantly, more complicated tools, such as DANN and Eigen, based on the combination of CADD’s inference model and deep learning, actually performed worse than CADD itself on our tests. Some may argue that this lack of precision is due to the fact that CADD, DANN and Eigen were developed with more attention paid to the variation in noncoding regions. Yet, that doesn’t explain why our own hybrid semi-supervised model, which was absolutely focused on the exome, didn’t beat its purely supervised sibling (though it did outperform most of the other scores we tested). We believe that a lot more research should be invested into unsupervised learning to uncover its full potential (or the lack thereof).

**Table 3.** Average precision score.

| Score | Test I | Test II |
| --- | --- | --- |
| CADD | 0.79 (0.70-0.87) | 0.60 (0.58-0.61) |
| DANN | 0.84 (0.77-0.90) | 0.55 (0.53-0.56) |
| Eigen | 0.81 (0.73-0.89) | 0.57 (0.56-0.58) |
| FATHMM | 0.83 (0.75-0.90) | 0.83 (0.82-0.84) |
| GERP++ | 0.77 (0.69-0.85) | 0.44 (0.43-0.45) |
| LRT | 0.87 (0.82-0.92) | 0.64 (0.63-0.65) |
| **MLP** | 0.91 (0.82-0.94) | 0.89 (0.88-0.91) |
| MetaLR | 0.88 (0.81-0.94) | 0.87 (0.87-0.89) |
| MetaSVM | 0.91 (0.87-0.95) | 0.87 (0.86-0.88) |
| MutationAssessor | 0.78 (0.70-0.85) | 0.67 (0.66-0.68) |
| MutationTaster | 0.91 (0.87-0.95) | 0.67 (0.67-0.68) |
| PROVEAN | 0.76 (0.67-0.85) | 0.61 (0.59-0.62) |
| PolyPhen HDIV | 0.80 (0.73-0.86) | 0.67 (0.66-0.68) |
| PolyPhen HVAR | 0.77 (0.69-0.84) | 0.66 (0.65-0.68) |
| SIFT | 0.79 (0.72-0.85) | 0.67 (0.66-0.68) |
| SiPhy 29-way | 0.75 (0.65-0.84) | 0.47 (0.45-0.48) |
| phastCons 100-way | 0.86 (0.81-0.90) | 0.66 (0.66-0.67) |
| phyloP 100-way | 0.89 (0.83-0.94) | 0.56 (0.54-0.57) |
| phyloP 7-way | 0.79 (0.70-0.86) | 0.46 (0.45-0.47) |
| **sdAE** | 0.92 (0.86-0.96) | 0.87 (0.86-0.87) |

## Conclusion

Here we successfully explored the possibility to efficiently utilise deep learning models to discriminate neutral and likely pathogenic nsSNVs. We tried to use two distinct architectures, one of which made use of unsupervised learning, and optimised hyper-parameters using a genetic algorithm. Although this work was not conceived as a tool comparison, but rather an exploratory study, our results proved that even relatively simple modern neural networks significantly improve prediction accuracy of a deleteriousness prediction tool. Though our semi-supervised model didn’t outperform its purely supervised sibling, it bested most of the scores we tested in the study. Our supervised model showed superior average accuracy as compared to other scores, especially other deep learning-based tools. We have created an open-access web-server so that other could easily use our best classifier: *http://score.generesearch.ru*.

## Supporting Information

**S1 File. Additional benchmarks.** Multiple additional performance indicators calculated on a range of binary cutoff thresholds.

